# A comprehensive analysis of RNA sequences reveals macroscopic somatic clonal expansion across normal tissues

**DOI:** 10.1101/416339

**Authors:** Keren Yizhak, Francois Aguet, Jaegil Kim, Julian Hess, Kirsten Kubler, Jonna Grimsby, Ruslana Frazer, Hailei Zhang, Nicholas J. Haradhvala, Daniel Rosebrock, Dimitri Livitz, Xiao Li, Eila-Arich Landkof, Noam Shoresh, Chip Stewart, Ayelet Segre, Philip A. Branton, Paz Polak, Kristin Ardlie, Gad Getz

## Abstract

Cancer genome studies have significantly advanced our knowledge of somatic mutations. However, how these mutations accumulate in normal cells and whether they promote pre-cancerous lesions remains poorly understood. Here we perform a comprehensive analysis of normal tissues by utilizing RNA sequencing data from ∼6,700 samples across 29 normal tissues collected as part of the Genotype-Tissue Expression (GTEx) project. We identify somatic mutations using a newly developed pipeline, RNA-MuTect, for calling somatic mutations directly from RNA-seq samples and their matched-normal DNA. When applied to the GTEx dataset, we detect multiple variants across different tissues and find that mutation burden is associated with both the age of the individual and tissue proliferation rate. We also detect hotspot cancer mutations that share tissue specificity with their matched cancer type. This study is the first to analyze a large number of samples across multiple normal tissues, identifying clones with genomic aberrations observed in cancer.

## Introduction

As cells divide during life they accumulate somatic mutations, most of which are thought to be either neutral or slightly deleterious (McFarland et al., 2013). However, a few may increase cellular fitness and contribute to clonal expansion. This process was found to be associated with aging as well as different diseases such as coronary heart disease (Jaiswal et al., 2014; Vermulst et al., 2008) and cancer. In cancer, several such events that accumulate over time can promote uncontrolled cellular growth. Indeed, international efforts such as The Cancer Genome Atlas (TCGA) and the International Cancer Genome Consortium (ICGC) have made a major contribution to our understanding of the molecular and cellular aspects of this disease (Abeshouse et al., 2015; Brennan et al., 2013; Burk et al., 2017; Genovese et al., 2017; Hoadley et al., 2018; Kim et al., 2017; Knijnenburg et al., 2018; Lin et al., 2014; Zheng et al., 2016). However, these studies provided only partial information regarding cancer initiation and progression. Acknowledging this gap, recent studies have focused on somatic mutations in normal human tissues and pre-cancerous lesions, aiming to identify early clonal expansions (Aghili et al., 2014; Beane et al., 2017; Blokzijl et al., 2016; Cai et al., 2014; Cooper et al., 2015; Genovese et al., 2014; Izumchenko et al., 2015; Jacobs et al., 2012; Jaiswal et al., 2014; Krimmel et al., 2016; M et al., 2015; Martincorena et al., 2015; McKerrell et al., 2015; Millikan et al., 1995; Nair et al., 2016; O’Huallachain et al., 2012; Ooi et al., 2014; Ståhl et al., 2011; Vijg, 2014; Vinayanuwattikun et al., 2016; Waki et al., 2017; Welch et al., 2012; Yadav et al., 2016). In blood, clonal expansion was detected, and was enriched with mutations in several genes that have been previously implicated in hematologic cancers. Importantly, it was shown that the existence of clones in the blood increased the risk for developing these types of cancers, as well as other diseases (Genovese et al., 2014; Jaiswal et al., 2014). In skin tissue, a work by Martincorena *et al.* (Martincorena et al., 2015) studied multiple needle biopsies of normal skin from 4 individuals, using ultra-deep sequencing of 74 cancer genes. A high burden of low allele frequency mutations was detected, including in genes that are frequently mutated in squamous cell carcinoma. Overall, these initial findings emphasize the need to comprehensively map the prevalence of clonal expansion across many human tissues and study their properties. Such findings may highlight early events in tumor development.

In this study we leveraged the massive collection of diverse normal tissues generated by the Genotype-Tissue Expression (GTEx) (Lonsdale et al., 2013) project. As part of GTEx, RNA-seq data were generated from over 30 primary tissues across hundreds of individuals, as well as whole genome and exome sequencing data of DNA extracted from matched blood samples (release V7; **Methods**). To study clonal expansion using this rich data we first developed a new pipeline, RNA-MuTect, for detecting somatic mutations from RNA rather than DNA sequencing data. Of note, previous studies utilizing RNA-seq for mutation calling have focused on germline variants (Piskol et al., 2013), or have used these data as an additional validation resource for mutations detected in the DNA (Radenbaugh et al., 2014). Here, we first validated RNA-MuTect using cancer and normal TCGA samples for which both DNA and RNA were extracted from the same samples, demonstrating its high sensitivity and precision. We then applied RNA-MuTect to identify somatic mutations across the entire GTEx dataset, and detected low allele frequency somatic mutations in different tissue types from many individuals.

## Results

### A pipeline for detecting somatic mutations using RNA-seq data

To develop a pipeline for detection of somatic mutations from RNA-seq data, and properly evaluate its performance, the correct set of mutations in the sample should be known. An example of such a dataset is a one in which DNA and RNA are simultaneously extracted from the same tissue portion. Therefore, we first collected a training set of 243 TCGA tumor samples, representing 6 tumor types, for which both DNA and RNA were co-isolated from the same tumor cell population (**Supplementary Table 1**). We started by identifying all the somatic point mutations that occur in these samples based on their tumor and patient-matched normal blood DNA whole exome sequencing data. This was done by applying MuTect (Cibulskis et al., 2013), a method for detecting somatic point mutations in cancer samples, as well as additional filtering criteria designed for the analysis of DNA sequencing data (**Methods**). Overall, we identified 75,388 point mutations across these 243 cases, ranging from a median of 75 to 414 mutations per sample across tumor types (**Figure 1A**). Next, for each of these cases, we detected somatic mutations by applying MuTect to the RNA-seq data from the tumor against the matched normal DNA, and applied the relevant DNA filtering tools to the RNA-based somatic mutations (**Methods**). The initial RNA analysis yielded a much larger number of single nucleotide variations (359,982), ranging from a median of 1075 to 1966 mutations per tumor type (**Figure 1A**). Comparison between the DNA- and RNA-based mutations showed significant discrepancies; 65% of the DNA-based mutations were not detected using the RNA, and 92% RNA-based mutations were not detected using the DNA analysis (**Figure 1B**).

**Figure 1:**
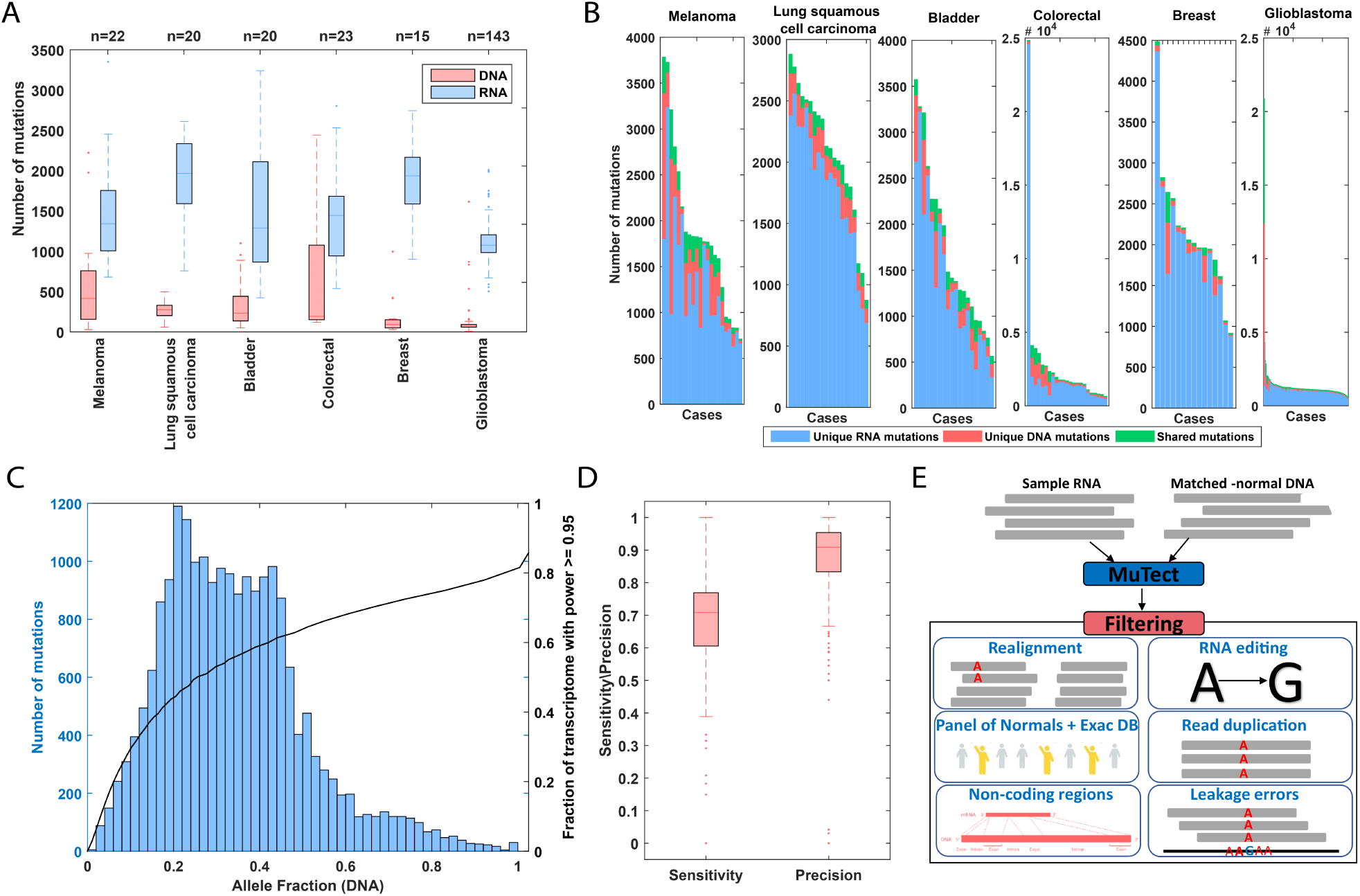
**(A)** Total number of mutations detected in DNA and RNA across samples in each TCGA cohort before filtering. **(B)** Number of mutations initially detected in both DNA and RNA (green), DNA alone (red) and RNA alone (blue), in samples from different cohorts. **(C)** Distribution of allele fraction in the DNA, of mutations detected in both DNA and RNA. The black line indicates the fraction of the transcriptome covered to a depth that provides ≥95% detection power. **(D)** Sensitivity and precision out of powered sites, across all studied cases. Box plots show median, 25^th^ and 75^th^ percentiles. The whiskers extend to the most extreme data points not considered outliers, and the outliers are plotted individually using the ‘.’ symbol. **(E)** A summary of the RNA-MuTect pipeline covering main filtering steps (**Methods**).

One obvious reason for not detecting mutations in the RNA is that the coverage may be insufficient in lowly expressed genes. We therefore performed a power analysis that examines which portion of the transcriptome is sequenced to a sufficient coverage that will allow us to detect mutations of different allele fraction (**Methods**). This analysis showed that in a typical sample, only 55% of the transcriptome had sufficient power (≥95%) to detect mutations at the median allele fraction of 0.32 (**Figure 1C**). When considering the actual allele fractions of the detected mutations and the coverage in the corresponding sample, we found that we are powered to detect only 33% of the mutations detected in the DNA. Out of the sites for which we had sufficient coverage (24,716/75,388), MuTect detected a much larger fraction of mutations, 82% (20,418) vs. 27% across all DNA-based mutations. We next examined the 4,298 mutations that were powered but not detected in the RNA. We found that for 3,198 of them (74%, an average of 13 mutations/sample) there was no or very low evidence for the alternate allele in the RNA (**Figure S1**). This could be a result of nonsense-mediated decay, expression coming from non-tumor cells and allele specific expression (**Figure S1**). Interestingly, the latter did not seem to have a major contribution as only 41 mutations showed such a pattern (**Methods, Supplementary Table 2**).

While the ability to detect mutations in RNA-seq data is clearly limited, a greater challenge is to filter out all the excessive mutations detected only in the RNA. To address this challenge we developed the RNA-MuTect pipeline that includes several filtering steps designed for RNA-seq data, such as removal of RNA editing sites, RNA-based sequencing errors, RNA-alignment artifacts and others (**Methods; Figure 1E**). When applied to the data, this pipeline filtered out the vast majority (93%; 334,345) of the RNA mutations. To estimate the false positive rate of the remaining mutations, we focused on RNA mutations with sufficient detection power in the DNA (22,687; 89%) and found that 90% (20,298) were indeed detected in the DNA (median precision of 0.91 across samples; **Figure 1D**), with a median of 3 mutations per sample that were detected in the RNA alone. The majority of the mutations detected only in the RNA (62%; 1484) were C>T mutations. A known RNA editing process involves the deamination of a cytidine base into a uridine base (Blanc and Davidson, 2003). These RNA editing alterations are less frequent than the more common A>G substitutions and constitute only 1% of the DARNED database (Kiran and Baranov, 2010), which we used for filtering. The observed rate may suggest that these mutations reflect yet undiscovered C>T editing events. However, we cannot rule out the possibility that these are false positives that were not filtered out by our pipeline. The massive filtering process required to achieve high precision, had only a slight effect on sensitivity, removing 2,511 (10%) mutations that were detected in the DNA and retaining a high overall median sensitivity of 0.7 (**Figure 1D**, **Figure S1**).

To evaluate the robustness of RNA-MuTect on an independent set, including in other tumor types, we collected an additional validation set of 303 TCGA samples representing 6 different tumor types (5 of which are different from our training set, **Supplementary Table 1**). Similarly, in these samples RNA and DNA were simultaneously co-isolated from the same biopsy portion. Applying RNA-MuTect to this set we were able to achieve high sensitivity and precision, having excellent agreement with the results on the training set (sensitivity of 0.72 and precision of 0.87, **Figure S2**). Similar to the training set, RNA-seq coverage provided sufficient power to detect only 31% of the mutations found in the DNA, out of which 72% were correctly detected in the RNA. On the other hand, 95% of the mutations originally detected by MuTect in the RNA are filtered out, and 87% of the powered RNA mutations are also detected in the DNA.

In this validation set we found a median of 4 mutations per sample that were detected only in the RNA. It is important to note that while we have decided to take a conservative approach in our validation, and consider all mutations detected in the RNA but not in the DNA as false positives, it is possible that, either (1) these are true mutations found only in the RNA and were generated via RNA-specific processes, or (2) the more likely scenario, that they exist in the DNA, but given their allele frequency and DNA coverage we were actually not powered to detect them in the DNA. Specifically, our power calculations assume that the mutation allele fractions are the same in the DNA and RNA, which is equivalent to assuming that the expression levels of a gene across different cells in the sample depend only on the gene’s local copy-number and not on the cell type. However, it is likely that genes have different expression levels in, for instance, tumor and stroma cells. To test whether the mutations found only in the RNA (in sites powered in the DNA) are real mutations, we examined the correlation between the number of RNA-specific mutations and the number of mutations detected in both RNA and DNA (i.e. the true positives). We indeed found a high correlation between these counts (Spearman R = 0.6, P-value = 4.2×10^−30^), suggesting that many of these are likely real, as we would not have expected a correlation between false positive (generated by noise) and true positive mutations. Nonetheless, to be highly conservative, throughout this study we consider powered mutations that were detected only in the RNA as false positives.

### *De novo* mutation calling from TCGA RNA-seq data reveals driver genes and mutational signatures

Next, we wanted to test whether we can discover cancer genes and mutational signatures based on the RNA-called mutations compared to the DNA-called ones in our training set. To this end we first applied MutSigCV (Lawrence et al., 2014), an algorithm for detecting significantly mutated genes, to both the DNA and RNA detected mutations separately. 15 driver genes were detected in this multi-cancer cohort based on DNA and 18 based on RNA mutation calls, with 10 genes overlapping between the two groups (**Figure 2A** and **S1**). Examining our ability to detect mutations in the remaining 5 genes that were detected only in the DNA, we found that indeed, we were only powered to detect these mutations in a subset of the samples, hence their evidence and significance dropped below MutSigCV’s threshold (**Figure 2B**). Overall, we found that the power to call RNA mutations in significantly mutated genes is significantly greater than the power across all DNA mutations (one-sided Wilcoxon P-value = 2.1×10^−47^; **Figure S1**). Of note, two out of five genes, *FGFR3* and *CDH1*, that were found to be significantly mutated only based on the RNA calls, are known cancer genes. The lower overall mutation frequency of the RNA-based mutations can increase the power to find cancer genes, if mutations in them are detected at a higher proportion than background mutations (potentially due to higher expression levels of cancer genes). We finally applied MutSigCV to our validation set and found that similarly, 10 out of 12 DNA-based significantly mutated genes were also considered significantly mutated based on RNA calls (**Figure S2**).

**Figure 2:**
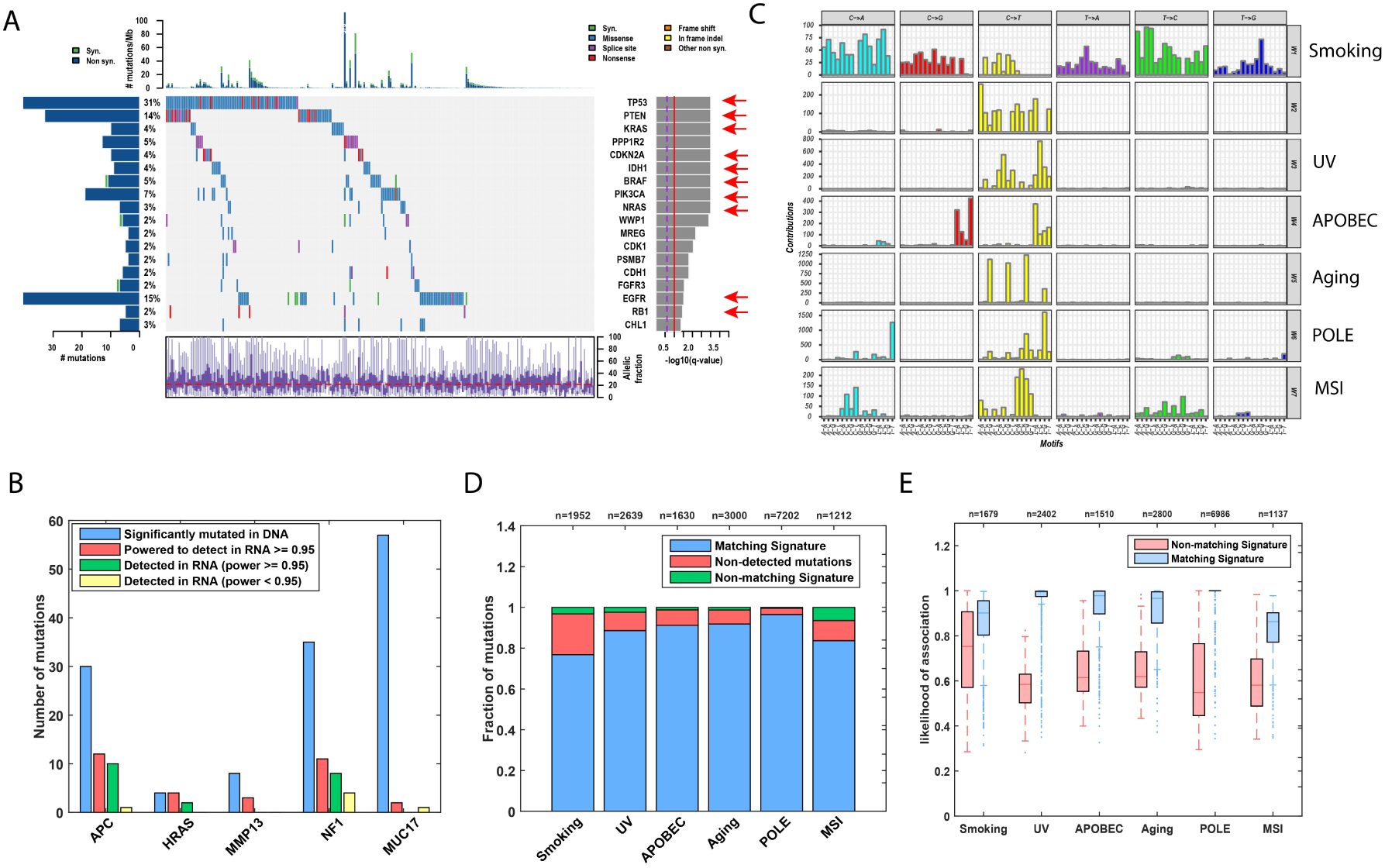
**(A)** CoMut plot derived from a MutSigCV analysis based on mutations detected in RNA. Identified cancer genes (q-value < 0.05) with their frequency, mutation type and allele fraction in the studied cohort are presented. Genes marked with a red arrow were identified as significantly mutated also in the DNA. **(B)** Significantly mutated genes identified by MutSigCV using the DNA-detected mutations. Blue: number of mutations detected in each gene in the DNA; red: number of mutations for each of these genes that are powered in the RNA; Green and yellow: number of mutations found in the RNA in powered and underpowered sites, respectively. **(C)** Mutational signatures identified based on mutations detected in the RNA. The mutational signatures identified are: a mixture of smoking and nucleotide-excision repair signatures (W1, COSMIC signatures 4 and 5, cosine similarities of 0.7 and 0.75, respectively); UV (W3, COSMIC signature 5, cosine similarity = 0.95); APOBEC (W4, COSMIC signature 13, cosine similarity = 0.9); Aging (W5, COSMIC signature 1, cosine similarity = 0.9); POLE (W6, COSMIC signature 10, cosine similarity = 0.88) and MSI (W7, COSMIC signature 15, cosine similarity = 0.8). **(D)** Fraction of RNA mutations associated with the same (blue) or a different (red) signature in the DNA, as well as undetected mutations in the DNA (green). **(E)** Likelihood of association of each RNA mutation associated with the same (blue) or a different (red) signature in the DNA.

We next examined which mutational signatures were active in the analyzed dataset using our SignatureAnalyzer which applies a Bayesian Non-negative matrix factorization approach to identify mutational signatures (Kim et al., 2016). Out of 8 mutational signatures discovered in the DNA (**Figure S1**), 6 were also discovered in the RNA, including Aging, UV, POLE, APOBEC, MSI and a mixture of Smoking and other signatures (**Figure 2C**). The majority of mutations associated with the two signatures undetected in the RNA (MSI and oxidative damage) originated from 4 samples (1 MSI colon cancer and 3 glioblastoma with oxidative damage), all of which had very low median allele fraction (0.029) and hence these mutations were largely missed in the RNA (**Figure S1**). When examining individual mutations while focusing on powered sites with a high assignment probability (**Methods**), we found that 87% of the mutations assigned to a specific mutational signature in the RNA, were assigned to the same mutational signature in the DNA (**Figure 2D**). Moreover, mutations that were not assigned to the same signatures had inconclusive assignments in the DNA (**Methods**; one-sided Wilcoxon P-value < 2.3×10^−13^, **Figure 2E**). Our analysis also discovered, in the RNA, a yet unreported mutational signature, which is dominated by C>T mutations, and of which the majority originated from a single colon cancer sample (**Figure 2C**). Of these mutations, 75% were powered but not detected in the DNA, suggesting that this signature may reflect the C>U RNA editing process mentioned above. When applying SignatureAnalyzer to our validation set we found that 4 out of 5 signatures detected based on the DNA calls were also detected based on the RNA calls (**Figure S2**). Overall, despite the apparent limitations in calling mutations *de novo* from RNA-seq data, we found that most cancer genes as well as mutational processes are revealed from these data alone.

### Validating RNA-MuTect in normal tissues

To validate RNA-MuTect in normal samples we collected 35 samples, from tumor-adjacent normal samples collected in TCGA, in which DNA and RNA were simultaneously co-isolated from the same tissue biopsy portion (**Supplementary Table 1**). Of note, these samples were found not to be contaminated with tumor cells (Taylor-Weiner et al., 2018). The average allele fraction of the mutations detected in these samples was 0.06, indicating that the discovered mutations are present in a small fraction of cells, as expected in normal tissues. Examining our ability to detect in the RNA mutations that were found in the DNA, we found that out of 8 mutations detected in the DNA that had sufficient coverage (power≥95%) to be detected in the RNA, 3 were indeed detected. Out of the remaining 5 mutations, 4 had no evidence in the RNA and one had low evidence of only 2 reads, therefore falling below our detection threshold (**Supplementary Table 3**). In addition, we examined how many of the mutations detected in the RNA had evidence in the DNA. First, similar to our finding in the DNA, the average allele fraction of the detected mutations was 0.07. Overall, we found that out of 86 RNA mutations that had sufficient coverage to be detected in the DNA, 13 mutations were indeed detected. The median and average number of mutations per sample that were detected only in the RNA, and therefore we consider as false positives (see above), are 1 and 2 sites, respectively (**Supplementary Table 3**). As only half of the mutations detected in the RNA had sufficient coverage to be detected in the DNA, we estimate that in correspondence with our findings above, the median number of false positive mutations per sample is 4. Overall, when applying RNA-MuTect to normal samples with proper co-extracted DNA and RNA data, we found that DNA mutations are detected in the RNA whenever reads supporting the mutations exist. More importantly, and in similar to our results in cancer samples, RNA-MuTect detected a low number of potential false positive calls per sample in normal tissues as well.

### Somatic mutations in normal tissues

After establishing RNA-MuTect’s performance on TCGA samples, we sought to study somatic mutations in normal tissues by analyzing RNA-seq data from the GTEx project. Notably, the identification of somatic mutations in normal tissues is challenging. Specifically, in order for mutations to be detected in a normal tissue, they need to exist in a large enough fraction of the RNA molecules which likely originate from different cells (**Figure 3A**). That is, a clone that consists of a sizable fraction of cells in the sample needs to exist, as well as share and express these somatic mutations. The mutation detection threshold would then depend on the clonal diversity of the sample, depth of sequencing and the expression level of the mutated gene. This is in contrast to tumor samples that contain large clones, where calling somatic mutations from RNA-seq is feasible, as demonstrated above. Additionally, bulk RNA-seq from normal tissues is comprised of a diversity of cell types, including muscle and fat cells that are not proliferating, and therefore are unlikely to contribute to an expanded clone. Furthermore, in this dataset, RNA was extracted from a relatively large amount of tissue material, limiting our ability to identify mutations present in small clonal populations. Overall, these parameters are expected to affect our precision level reflecting the fewer true positive mutations. Despite these limitations, we applied RNA-MuTect to 6,707 RNA-seq samples against their matched-blood DNA, spanning 29 human tissues across 488 individuals (**Methods**), and detected 8,870 somatic mutations in 37% (2,519) of the samples representing nearly all individuals (95%, 467/488; **Figure 3B**, **S3 and Supplementary Table 4**). The median allele fraction of the mutations was 0.05, indicating that the discovered mutations are present in a small fraction of cells (**Figure 3C**). Given our estimation indicating a median of 4 false positives mutations per sample, we still find that 375 samples have more than 4 mutations (i.e., at a confidence level of 50%), and that 106 samples have more than 13 mutations (equivalent to a confidence level of 80%, **Figure 3B**). However, it is important to note that mutations detected in samples with an overall low number of mutations, may still truly exist in the sample (as discussed above), even though the overall number is below the false positive level. For example, known cancer mutations that can increase the fitness of the cells have a higher prior probability of being real, as will be further discussed below. Overall, the results above indicate that macroscopic clones exist in different normal tissue types.

**Figure 3:**
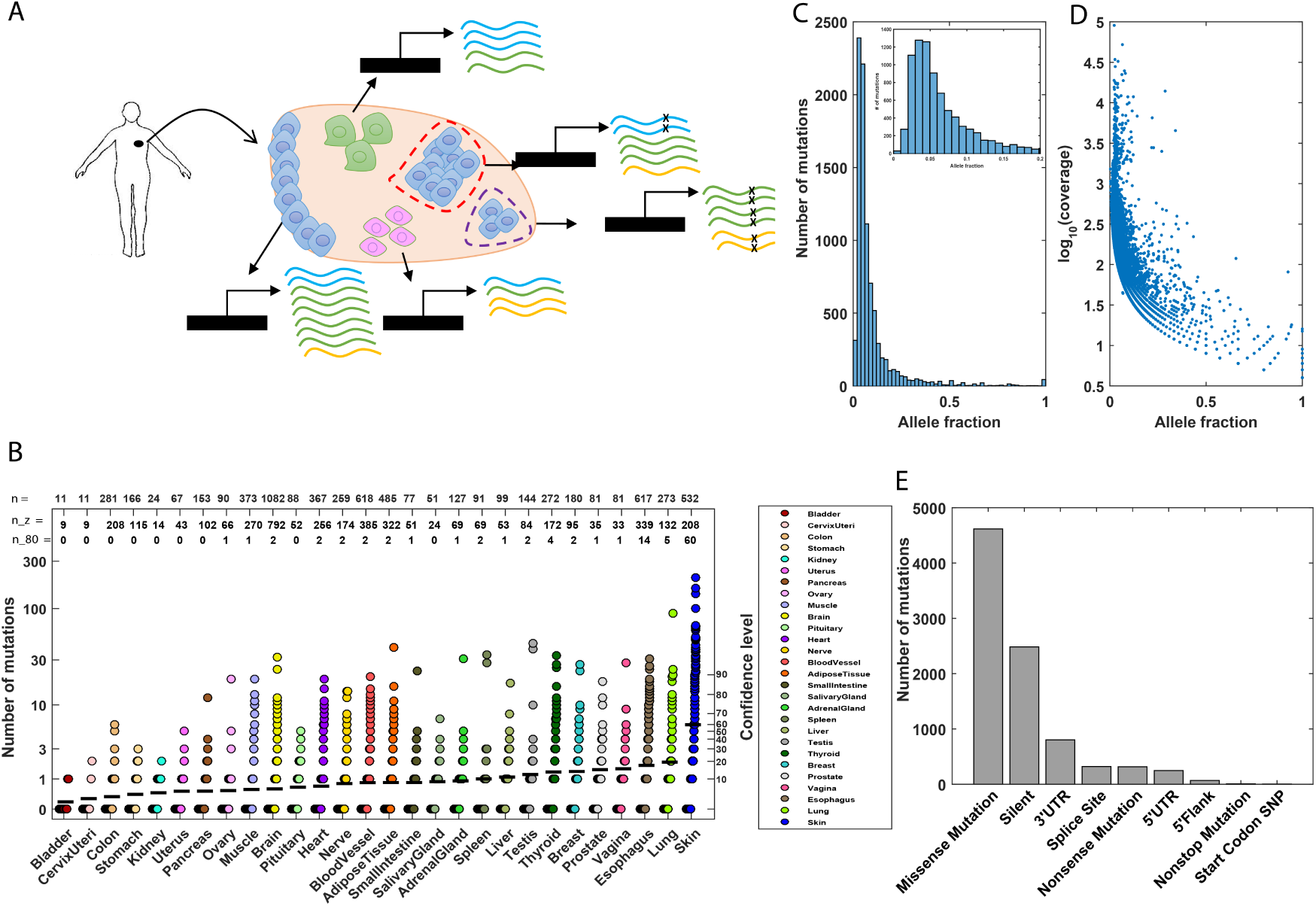
**(A)** An illustration of the composition of bulk RNA extracted from a normal human tissue. The biopsy consists of three different cell types that express different transcripts (marked in blue, green and yellow) at different levels. Blue cells represent cells with a higher probability to form clones. Two clones are shown, a small and a large one, marked in purple and red, respectively. Mutated reads are marked with an x. The allele fractions of the mutations in the blue and green genes are the same (0.25; 2/8 and 4/16 reads, respectively) despite the fact that the clones are of different size. Additionally, the allele fraction of the mutation in the yellow gene is higher than the allele fraction of the mutations in the blue and green genes (0.33; 2/6 reads), despite the fact that the yellow mutation is supported by the same, or a smaller number of reads. These scenarios illustrate the difficulty in identifying somatic mutations in a bulk normal tissue due to mixture of cell types and the small size clones. Moreover, inferring clone size is limited due to different cell types that exist in different proportions and express transcripts at different levels. **(B)** Number of mutations detected in the RNA-seq of all studied tissues. Each sample is represented with a circle. The black horizontal line represents the mean number of mutations in each tissue type. A confidence level based on our estimation of false positives in the validation data is indicated in the right y-axis. Specifically, this confidence level is computed as the x^th^ percentile on the number of false positive calls (RNA only mutations in DNA powered sites) found in the validation set. n, n_z and n_80, indicate: (1) the total number of samples analyzed in each tissue; (2) the number of samples in which no mutations were detected; and (3) the number of samples in which more than 13 mutations were found (equivalent to a confidence level of 80%), respectively. **(C)** Distribution of allele fraction across all samples in which somatic mutations were detected. Inset: mutations with allele fraction ≤ 0.2. **(D)** Allele fraction as a function of the log_10_(coverage) for all detected mutations. **(E)** Summary of the mutation types detected in the data.

As expected, owing to the higher sensitivity to detect low allele fraction mutation in highly covered sites, a strong negative correlation between sequencing coverage and allele fraction was found (Spearman R = −0.8, P-value < 1^-200^, **Figure 3D**). After correcting for the power to detect mutations (given the mutation allele fraction and the effective gene coverage; **Methods**), we observed a negative relation between expression level and expected number of mutations (**Methods, Figure S4**). Similar findings suggesting transcription-coupled repair were observed in cancer (Chapman et al., 2011; Lawrence et al., 2013; Pleasance et al., 2010), thus further supporting our data. The majority of the detected mutations were missense mutations (4616, 52%), similar to their frequency in TCGA samples (1,540,318; 53%, based on the TCGA consensus somatic mutation call set, MC3, doi:10.7303/syn7214402, **Figure 3E**). An additional comparison to TCGA data identified 137 mutations in normal tissues that were exactly the same as mutations found in the corresponding cancer type (**Supplementary Table 5 and Figure S5**).

In the GTEx study, and as opposed to TCGA data used here for training and validation, DNA and RNA were extracted from adjacent but different samples. Considering the low allele frequency of the detected mutations, and the fact that the adjacent biopsy may have a different composition of these different cells, the odds of identifying these microscopic clones in the adjacent tissue extremely low and unpredictable. Nevertheless, we attempted to validate 28 mutations by performing deep sequencing of the DNA at the corresponding sites. Although we did not expect to find any of the mutations from these small clones in adjacent tissue, we were able to validate in the DNA (with >=30 reads) 5 of the 28 mutations that were attempted (**Supplementary Table 6**; **Methods**). In addition, below we provide multiple lines of evidence that indicate that many of the detected mutations are true somatic mutations.

We found that the tissues harboring the greatest number of mutations are skin, lung and esophagus, all of which are associated with environmental carcinogenic factors such as UV radiation, smoking and nutritional habits (Beate et al., 2011; Gilchrest et al., 1999; Kamangar et al., 2009). Importantly, arranging tissues by their median coverage results with a completely different order, suggesting that coverage had no major effect on this finding (**Figure S3, Supplementary Table 7**). Looking at tissue sub-regions, we found that sun-exposed skin samples have the highest number of mutations, and esophagus mucosa, from which esophageal squamous cell carcinoma are derived, is the second-highest (**Figure S3**). The only tissue with a significant difference in number of mutations between males and females was breast (two-sided Wilcoxon P-value = 2.1×10^−5^, **Figure S3**). This reflects the observation that breast tissues sample from males in the GTEx dataset are mainly composed of fat cells while female breast tissues are composed of epithelial cells as well.

Several factors can affect the number of mutations accumulated in normal tissues. These include the age of the individual, the accumulated DNA damage and the tissue’s propensity for forming macroscopic clones. All are expected to be more prominent in tissues with a higher cell proliferation rate (Tomasetti et al., 2013). To test these associations in our data we first examined the relation between the age of the individual and the average number of accumulated mutations across tissues. We found a significant increase after the age of 45 in both the average number of aging mutations (CpG>T) as well as in the average number of all mutations (one-sided Wilcoxon test P-value = 0.001 and P-value = 2.2×10^−4^, respectively, **Figure 4A**, top panels). Importantly, a significant association was observed even after controlling for the number of tissues sequenced in each individual (**Supplementary Table 8**), and when splitting all individuals to three age groups (**Figure S6**). As expected, when considering tissues with a higher level of cell proliferation (as determined by the expression level of *MKI67*, a marker of proliferation, **Methods**, **Supplementary Table 9**), this relationship becomes more significant for the total number of mutations (P-value = 2.2×10^−5^), and remains similar for the aging mutations (P-value = 0.003). Exploring the number of mutations only in tissues with a lower cell proliferation, the association with age was not significant (P-value = 0.59). Next, we tested the association with age in each tissue separately. A significant association was detected only in skin and esophagus tissues (P-value = 2.1×10^−6^ and P-value = 1.5×10^−5^, respectively, **Figure 4A**, bottom panels). When considering sun-exposed and non sun-exposed skin separately, we found that in both cases the number of mutations observed increase with age, but significantly more in sun-exposed skin (P-value = 3.9×10^−8^ and P-value = 0.04 for sun-exposed and non-exposed, respectively, **Figure S6**). Since these samples come from the same tissue type, and hence are likely to have a similar cell proliferation rate, this result emphasizes the environmental contribution to the accumulated damage. Similarly, when testing the different parts of the esophagous tissue (mucosa, gastroesophageal junction and muscularis), both the mucosa and the muscularis showed a significant association with age, though to a different extent (P-value = 8.4×10^−9^ and P-value = 0.02, respectively; **Figure S6**). In this case, while both the mucosa and the gastroesophageal junction are exposed to similar environmental stresses, the mucosa has a higher cell proliferation rate. The lack of association in other tissues can be a result of either low cell proliferation rates, or clones that were below our detection threshold. Finally, we directly examined the association between the expression level of *MKI67* across tissues and the number of accumulated mutations. We found a significantly higher expression level of *MKI67* in tissues with higher number of mutations (P-value = 8.2×10^−4^ and P-value = 1.2×10^−4^ for all primary and sub-region tissues, respectively, **Figure 4B and S6**, **Methods**). Overall, we found that in normal tissues, both age and exposure to mutagenic factors contribute to the number of accumulated mutations in tissues with a high cell proliferation rates (Alexandrov et al., 2015; Tomasetti et al., 2013).

**Figure 4:**
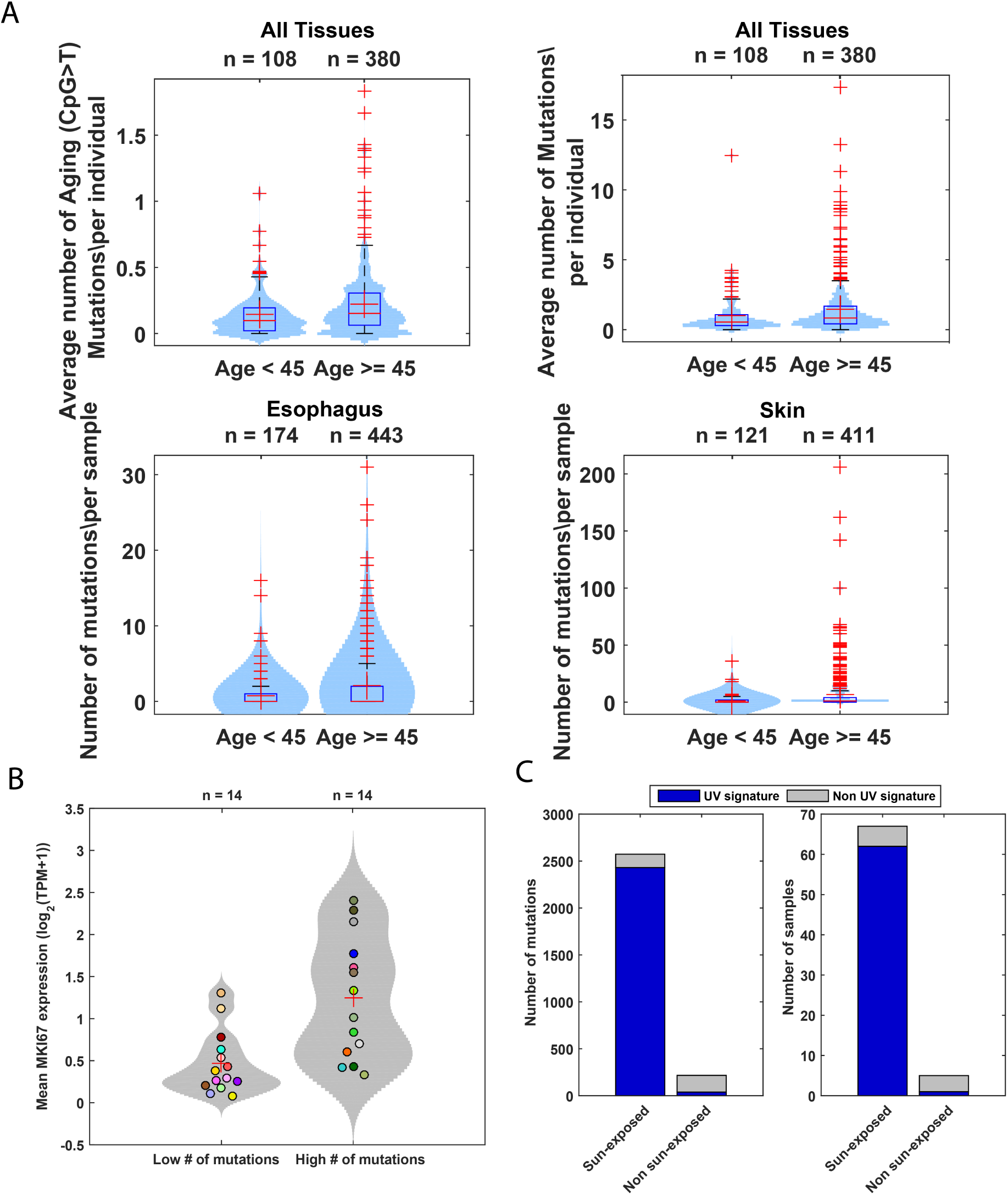
(A, top panels) Differences in the average number of aging-related and total number of mutations before and after the age of 45 (left and right panels, respectively); **(A, bottom panels)** Differences in mutation number in esophagus and skin samples before and after the age of 45 (left and right panel, respectively). Box plots show median, 25^th^ and 75 percentiles in each group. Red crosses represent the outliers and black crosses represent the mean **(B)** Difference in the expression of *MKI67* in tissues with lower and higher number of mutations than the median tissue. Dots: average expression per tissue type, color-coded according to Figure 3B. The average expression level of *MKI67* in each group is marked with a black cross, and the average expression level per tissue type is marked with a circle. **(C, left panel)** Number of mutations associated with the UV signature in sun-exposed and non sun-exposed skin samples; **(C, right panel)** number of samples associated with the UV signature in sun-exposed and non sun-exposed skin samples.

Going beyond the sample level, we next examined whether there are individuals with a significantly higher or lower number of mutations across their tissues, while correcting for their age, gender and race (**Methods**). We identified 10 individuals with a significantly higher number of mutations and 5 individuals with a significantly lower number of mutations (q-value < 0.1). In addition, we found that the 5 individuals with the lower number of mutations are significantly younger (two-sided Wilcoxon P-value = 5×10^−3^, with 4 individuals younger and one older than the median age of 45). No association with age was found for those with the higher number of mutations. It is important to note that there are likely additional factors that were not accounted for, such as the purity of the tissue sample (i.e. the fraction of cells in a tissue sample that belong to the specific tissue). Therefore, we cannot conclude that these individuals indeed have an intrinsic higher propensity to accumulate mutations.

Further examining our data, we identified a vagina sample taken from a 21-year-old female, which had 28 detected mutations, representing an outlier in this tissue (**Figure S7**). We suspected that this might have resulted from an infection with human papillomavirus (HPV), known to have a high prevalence in young women, although we did not find evidence of HPV in the RNA-seq. Indeed, the set of genes that were highly expressed in this sample were significantly enriched with immune-related genes (q-value = 5.09×10^−10^, **Supplementary Table 10**, **Methods**), and specifically included various immunoglobulins. B cells, which are responsible for virus neutralization, secrete immunoglobulins. A systemic humoral immune response was shown to be present in more than 60% of women infected with HPV in the genital tract (Carter et al., 1996). Additionally, the genes *APOBEC3A* and *APOBEC3B* were among the most highly expressed genes in this sample with a 3-fold increase over their tissue average. *APOBEC3* cytosine-deaminase enzymes are known to be active in HPV-infected tissues (Vartanian, 2008) and have a unique mutational signature. Consistently, 26 out of the 28 mutations in this sample were C>T mutations in the T<u>C</u>W motif, which are the canonical APOBEC mutations (Roberts et al., 2013) (**Figure S7**).

We next sought to identify mutational processes that are active in normal samples by applying SignatureAnalyzer (**Methods**). Since most samples have a small number of mutations, we analyzed only the 169 samples with ≥10 mutations. SignatureAnalyzer identified a single signature, the UV signature, in skin samples. The UV signature is common in melanoma and was recently reported for skin fibroblasts and normal skin samples (Martincorena et al., 2015; Saini et al., 2016). When examining sun-exposed and non-exposed skin separately, we found that the UV signature is active in 62 out of the 67 sun-exposed samples, and only in 1 of the 5 non sun-exposed samples (**Figure 4C**, Fisher P-value = 5.7×10^−4^). Interestingly, all the skin samples with ≥10 mutations analyzed here were from individuals of European ancestry, while none of the samples from individuals of African-ancestry analyzed in this study had more than 6 mutations, regardless of sun-exposure (one-sided Wilcoxon P-value = 0.84). Overall, skin was the only tissue that showed a significant difference between the total number of mutations detected in European vs. African-ancestry samples (one-sided Wilcoxon P-value = 1. 9×10^−5^, **Figure S3**). Of note, as most of the samples in this study (95%) have less than 10 mutations, the power to detect additional signatures is very low. To demonstrate this issue we used the TCGA data analyzed here in which 7 different mutational signatures were detected (**Figure 2C**). We selected a subset of the called mutations, such that the distribution of mutation number per sample is identical to that found in GTEx (**Methods**). Running SignatureAnalyzer on these data yielded only two signatures, the UV signature and an additional one most closely related to the aging signature but with a relatively low cosine similarity (**Figure S8**).

### Mutations in cancer genes in normal tissues

To determine whether somatic mutations in normal tissues occur in known cancer genes, we first tested the frequency of non-silent mutations in Cancer Gene Census (CGC) genes (Forbes et al., 2015). This set represents genes where mutations have been causally implicated in cancer. We found that 3% of the samples and 33% of the individuals carried at least one non-silent mutation in a CGC gene. To further investigate this finding we examined which tissues are significantly enriched with these non-silent mutations (**Methods**). Five tissues (skin, esophagus, adipose tissue, adrenal gland and uterus) were found to be significantly enriched with CGC genes (q-value < 0.05), while controlling for both gene length and coverage (**Figure S9**, **Methods**).

The most frequently mutated cancer gene in our data was *TP53*. Examining whether a significant difference in the number of mutations is found between samples carrying *TP53* mutations and those that do not, we found a significant increase in the *TP53*-associated samples (two-sided Wilcoxon P-value = 9.2×10^−9^). To test if these mutations provided a growth advantage to the cell, we checked whether they existed at a higher allele fraction relative to all other detected mutations in the same sample, focusing on samples with at least 3 mutations (**Methods**). Indeed, we found that the allele fraction of 12 out of 19 *TP53* mutations was in the top 2 quintiles, i.e. above the 60^th^ percentile (empirical P-value < 0.02, 6 fell above the top 80^th^ percentile, **Figure S9**). Of note, this observation is independent of *TP53* coverage, as for 18 out of 22 mutations, TP53 average coverage is found below the 60^th^ percentile as compared to other mutated genes detected in the sample (**Figure S9**). The second most mutated cancer gene in our data was *NOTCH1*, found in normal skin and esophagus, with a similar distribution of mutation types as previously reported in skin (Martincorena et al., 2015) (two-sided Wilcoxon P-value = 0.6, **Figure S9**).

We next examined whether any of ∼1760 recurrent cancer mutations are observed in normal tissues (**Supplementary Table 11**, **Methods**). We found 30 mutations in 8 tissues, spanning 27 hotspots in 12 genes (**Figure 5A, Supplementary Table 12**). The gene with the greatest number of detected hotspot mutations was *TP53*. Here we found a total of 16 *TP53* hotspot mutations in both skin and esophagus samples, with 14 of them being unique (**Figure 5A**). In total, 10 of these mutations were previously reported in either normal human skin, peritoneal or uterine lavage fluids taken from healthy women, or in human pluripotent stem cells (Krimmel et al., 2016; Martincorena et al., 2015; Merkle et al., 2017; Nair et al., 2016). Reviewing the IARC TP53 database (Petitjean et al., 2007), we found that all of these mutations were annotated as deleterious by SIFT (Kumar et al., 2009). Interestingly, although all of the mutations were annotated as loss-of-function in yeast, 3 of them, R248Q, R248W and R282W were reported to have gain of function activities (Muller and Vousden, 2014). R248Q knock-in mice showed an earlier onset of tumor formation and reduced lifespan, as well as an expansion of hematopoietic and mesenchymal stem cell progenitors (Hanel et al., 2013). The R248W variant was found to be involved in multiple gain of function activities, including promotion of cell invasion (Muller et al., 2009), increase in cell proliferation (Yan and Chen, 2009) and more (Muller and Vousden, 2014). The R282W variant was shown to increase colony formation (Scian et al., 2004). We found that these 14 mutations share some tissue specificity with the corresponding primary cancerous tissue, where 4 skin and 5 esophagus mutations were also observed in melanoma and esophagus TCGA samples, respectively (**Figure 5B**).

**Figure 5:**
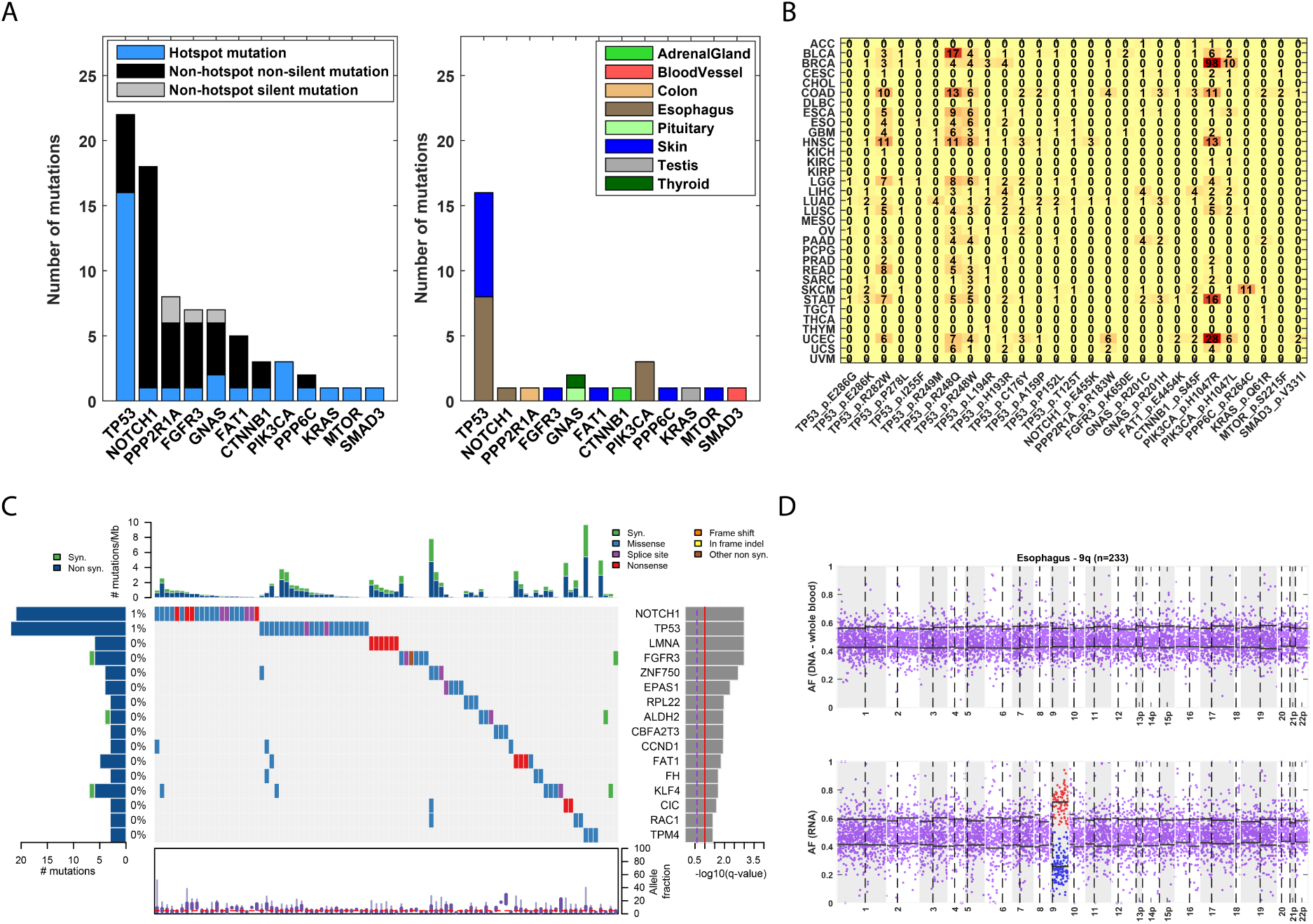
**(A)** Genes in which hotspot mutations were detected. Left panel: number of hotspot mutations detected in each gene, and number of silent and non-silent mutations that are not in hotspots. Right panel: normal tissues in which the hotspot mutations were detected. **(B)** Occurences of each hotspot mutation found in different TCGA cohorts. **(C)** CoMut plot for genes found to be significantly mutated in a pan-normal analysis, ordered by their significance level, showing 93 of 6707 samples with at least one mutation in these genes and the overall frequency among samples with at least one mutation. The distribution of allele fraction of mutations appears at the bottom. **(D)** Allelic imbalance in chromosme 9q of a normal esophagus sample. Top panel: allele fraction of heterozygous sites based on DNA from a matched-blood sample. Bottom panel: allele fraction of heterozygous sites based on RNA from the esophagus sample. The black horizonal lines indicate the mean allele fraction per chromosomal arm of sites with allele fraction smaller or greater than 0.5.

Considering the other 14 non-TP53 hotspot mutations, we found that all of them but two (FAT1 p.E4454K; FGFR3 p.K650E) are annotated as pathogenic by FATHMM (Shihab et al., 2013), and 7 share tissue specificity with the corresponding cancerous tissues (**Figure 5B**). Specifically, a hotspot mutation in *PPP2R1A*, a regulator of cell growth, which we detected in a normal colon sample, was also detected in colorectal cancer. While it is the β isoform (*PPP2R1B*) that was first discovered as a tumor suppressor in colon cancer cell-lines and primary tumors (Wang et al., 1998), this α isoform was later observed in a larger cohort of primary colon tumors (Muzny et al., 2012). The hotspot mutation in *CTNNB1* (β-catenin), found in the adrenal gland sample of a 58-year-old female, was previously detected in adrenocortical adenomas, as well as adrenocortical tumors (Bonnet et al., 2011; Leal et al., 2011; Zheng et al., 2016) resulting with Wnt/β-catenin pathway deregulation. *CTNNB1* was also found to be significantly mutated in the latter (Zheng et al., 2016), therefore possibly driving the disease. The *PPP6C* hotspot mutation detected in normal skin was identified in melanoma samples and similarly was found to be significantly mutated. Two *PIK3CA* mutations, P.1047L and p.1047R, that are common in multiple cancers including esophageal, are found here in normal esophagus mucosa samples. Lastly, the *KRAS* hotspot mutation (p.Q61R) found in a normal testis sample of a 58-year-old male, was previously detected in a testicular germ cell cancer. Examining whether these mutations had a higher allele fraction relative to all other detected mutations of that sample. Indeed, we found that the allele fraction of 14 out of 22 tested hotspot mutations fall above the 60th percentile, even when controlling for gene coverage (empirical P-value < 0.04, 9 fell above the top 80th percentile; **Methods** and **Figure S9**).

To further explore whether clonal expansion observed in normal tissues is in part due to positive selection of cancer genes, we computed a dN/dS ratio per gene, while taking into account the trinucleotide context and the mutational spectrum (**Methods**). We then examined whether an excess of non-silent mutations in cancer genes is detected in our data. Indeed, we found that both CGC genes as well as cancer significantly mutated genes (Lawrence et al., 2014) have a higher rate of non-silent mutations (one-sided Wilcoxon P-value = 3.4×10^−4^ and P-value = 9.3×10^−4^, respectively), suggesting that some of the mutations in them may confer a selective advantage.

To more specifically identify which of these cancer genes are significantly mutated, we performed a pan-normal analysis by applying MutSigCV (Lawrence et al., 2014) to all samples in which we detected at least one mutation while testing 718 known cancer genes (**Methods**, **Supplementary Table 13**). This analysis yielded 16 significantly mutated genes, with mutations spanning 17 tissues, 93 samples and 82 individuals (**Figure 5C and S9**). The two tissues with the greatest number of mutations in significantly mutated genes are the skin and esophagus, with 57 and 36 mutations, respectively. In addition to *TP53, NOTCH1* and *FAT1* that were previously detected as significantly mutated in normal skin (Martincorena et al., 2015), we also identified genes such as *RAC1* and *ZNF750*, both of which were reported as significantly mutated in melanoma and esophagus squamous cell carcinoma, respectively (Lawrence et al., 2014; Lin et al., 2014). Examining the allele fraction of the 79 non-silent mutations in these 16 genes, we found that 27 (35%) of them fall above the 80^th^ percentile, even when controlling for gene coverage (empirical P-value < 0.002, **Methods** and **Figure S9**).

Going beyond solid tissues, we examined whether somatic mutations are detected in the blood of the studied individuals (**Methods**). Focusing on a previously defined set of 332 single nucleotide variants detected in the blood of healthy individuals (Jaiswal et al., 2014), we identified 85 mutations across 82 individuals (17% of the studied individuals). Next, for each of the 82 individuals, we tested whether we can find the exact variant(s) in other solid tissues from the same individual. Only 7 mutations had evidence in at least one read in other tissues (each in a different individual), across different tissue types (5 brain, 1 thyroid and 1 heart, **Supplementary Table 14**). This result may represent the fact that these tissues contained some blood cells. In a previous study (Jaiswal et al., 2014), the authors found an increase in the number of detected mutations above the age of 70. Although the oldest person in our dataset is 70 years old, we did observe that these individuals had a significant trend towards being older (one-sided Wilcoxon P-value = 0.049).

### Allelic imbalance in normal tissues

We finally developed a method for identifying allelic imbalance events of chromosome arms, using RNA-seq data (**Methods**). We identified 8 esophagus mucosa samples that had allelic imbalance in 9q (**Figure 5D and S10**). Two out of the 8 samples also had a nonsense or missense mutation in *NOTCH1* (hypergeometric P-value = 0.02) which is located on the same chromosome arm. The allele fraction of these mutations was relatively high 0.22 and 0.12) and found at the top 20^th^ percentile of mutations in the corresponding samples. This may suggest that either the wild-type copy of these chromosome arms was lost, or that the mutated copy was gained. A frequent genomic amplifications in this gene was recently reported for esophageal squamous cell carcinoma (Kim et al., 2017). Interestingly, 9q loss was found to be more common in esophageal dysplasia than in esophageal squamous cell carcinoma (Shi et al., 2013). Its detection here in non-dysplastic lesions suggests that this may be an early event in the development of dysplasia. In similar to previous work studying normal skin samples (Martincorena et al., 2015), we found that both point mutations and allelic imbalance events in *NOTCH1* exist in normal esophagus. One additional sample carried mutations in both *TP53* and *FAT1*. An allelic imbalance in 22p and a mutation in *NOTCH1* was identified in an additional esophagus sample (**Figure S10**). Finally, we identified a testis sample with a strong allelic imbalance in 17p in which no point mutation was detected (**Figure S10**).

## Discussion

Here we studied clonality in normal tissues by leveraging the large collection of RNA-seq data taken from normal tissues as part of the GTEx project. To address this challenge we have first developed a pipeline for detecting somatic mutations from RNA-seq data. As expected, we found that many mutations do not have sufficient coverage in the RNA to be detected. However, the majority of those who are sufficiently covered, are detected using a DNA mutation calling pipeline (Cibulskis et al., 2013). On the other hand, we found that the RNA analysis also detected many RNA mutations that were not found in the DNA, the majority of them likely represent false positive calls. With this in mind we developed the RNA-MuTect pipeline. By applying a series of filtering criteria we were able to filter out the vast majority of these false calls and achieve overall high sensitivity and precision levels. The remaining mutations that were not filtered out and appear only in the RNA (a median of ∼4 per sample) are of particular interest. Although those mutations may simply represent false calls that were not filtered out by our approach, they might also be a result of different RNA editing processes that affect a macroscopic fraction of cells. Exploring these calls across all TCGA samples is out of the scope of this study, but can potentially reveal additional RNA editing events in cancer (Han et al., 2015; Paz-Yaacov et al., 2015).

Applying the RNA-MuTect pipeline to the GTEx dataset, we were able to start addressing the challenge of clonality in normal tissues on a large-scale, exploring thousands of samples across multiple tissues, covering individuals of different age, gender and race. Despite many potential limitations, including the expected frequency of clones within a given biopsy and the mixture of different cell types, we were able to detect many somatic mutations, some shared characteristics with mutations found in cancer and pre-cancer lesions. This includes cancer hotspot mutations, enrichment for non-silent mutations in cancer genes, allelic imbalance events, as well as association between age and the number of accumulated mutations. The two tissues in which we found the greatest number of cancer-related events were sun-exposed skin, which is associated with UV-damage (Gilchrest et al., 1999), and the esophagus mucosa, which is exposed to various environmental agents known to increase the risk for esophageal cancer (Kamangar et al., 2009). Both of these tissues also have a high cell proliferation rate.

Understanding the earliest genetic events that occur in human tissues may advance our understanding of aging and cancer. In cancer, studying early events is of great importance in aiding cancer detection and prevention. Studies characterizing the premalignant landscape of somatic mutations across normal tissues and blood cells are expected to transform our understanding of cancer biology, and help defining the sequencing of events leading to malignancy. Therefore, initiatives such as the Pre-Cancer Genome Atlas (Campbell et al., 2016) will significantly aid our ability to detect and treat the disease in its early stages. As all individuals in this study are deceased, we were unable to determine whether the detected clones would have eventually develop into cancer, upon acquisition of additional genetic abnormalities. However, studying samples longitudinally as they progress from a normal tissue to a pre-malignant lesion, that may then progress to cancer or remain at the pre-malignant state, will shed light on the combination and sequence of molecular events required for malignant transformation.

Finally, the overall low rate of cancer related events in our data (<10%), most likely reflects both our sensitivity and the fact that we have analyzed only a single biopsy from each tissue type in each individual. Given previous studies performing deep sequencing on much smaller tissue biopsies (Martincorena et al., 2015) it is reasonable to assume that had we analyzed many more biopsies from any given tissue type, and if those biopsies would have been more enriched with epithelial cells, a much larger number of somatic mutations across all normal tissues would have been detected, having additional overlaps with known cancer mutations. Certainly, not all these clones would develop into cancer, implying that while these clones have expanded to the point that they are detectable, they may remain harmless until, and only if, additional transforming events will be acquired in them. Early detection of cancer is a major effort in current times, as it may enable detection of the disease at a stage where it can be completely cured. However, as we presented here, some normal tissues harbor somatic mutations and have local clonal expansions. Therefore, early detection efforts will need to take this into consideration when approaching this challenge.

## Methods

### Mutation calling pipeline

TCGA DNA BAM files aligned to the NCBI Human Reference Genome Build GRCh37 (hg19), and RNA FASTQ files were downloaded from the Genomic Data Commons database (**Supplementary Table 1**). Sample contamination by DNA originating from a different individual was assessed using ContEst (Cibulskis et al., 2011). Somatic single nucleotide variations (sSNVs) were then detected using MuTect (Cibulskis et al., 2013). Following this standard procedure, we filtered sSNVs by: (1) removing potential DNA oxidation artifacts (Costello et al., 2013); (2) realigning identified sSNVs with NovoAlign (www.novocraft.com) and performing an additional iteration of MuTect with the newly aligned BAM files; (3) removing technology- and site-specific artifacts using a panel of ∼7000 TCGA normal samples (PoN filtering, see (Ellrott et al., 2018)). Finally, sSNVs were annotated using Oncotator (Ramos et al., 2015). RNA-seq data was aligned to the NCBI Human Reference Genome Build GRCh37 (hg19) using STAR (Dobin et al., 2013) (**Supplementary Table 15**). For initial calling of mutations from RNA-seq data we applied MuTect followed by the DNA-based PoN filtering. The other two filtering steps (oxoG and NovoAlign re-alignment) were not initially designed for RNA-seq data.

### Power analysis

Given a mutation with alternate allele count of *x* and reference allele count of *y* in a specific data type (DNA or RNA), we computed the power to detect it given a coverage *N* in the other data type (RNA or DNA, respectively). This was done by applying a beta-binomial model for observing at least *k* reads: 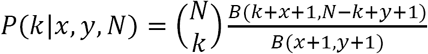 where *B* is the Beta function. To determine the minimal number of reads *k*, we first computed the error rate at the variant site, *r*, using the matched normal sample by taking the maximal allele fraction of the three possible alternate alleles and applying the Laplace correction with α=1. We then identified *k* as the number of alternate reads that have a probability <1% to be generated by the noise.

### The RNA-MuTect pipeline

The RNA-MuTect pipeline includes the following steps: (1) Applying MuTect to STAR-aligned RNA-seq BAMs with the ALLOW_N_CIGAR_READS flag and considering only mutations supported by ≥3 or ≥4 reads (for TCGA and GTEx analysis, respectively); (2) Applying the DNA PoN filtering as mentioned above; (3) A realignment filter for RNA-seq data where all reads aligned that span a candidate variant position from both the tumor (case) and normal (control) samples are realigned using HISAT2 (Kim et al., 2015) (Supplementary Table 14). We then performed an additional iteration of MuTect with the newly aligned BAM files; Only mutations that are kept by MuTect using both the STAR-aligned and HISAT-aligned BAM are kept for the next step; (4) Applying an RNA-seq PoN built based on a panel of ∼6500 GTEx samples, similar to the DNA PoN filter with a threshold score of −3 (Ellrott et al., 2018). The RNA-based PoN accounts for various RNA potential and recurrent artifacts, such as errors caused by reverse transcription of RNA to cDNA, RNA modifications and more. In addition, we removed 8 known hotspot mutations in *TP53* and *PIK3CA* from the PoN filter; as these are known cancer mutations, we had a much stronger prior that they are somatic events rather than technical artifacts (**Supplementary Table 16**); (5) Removing sites that were found in the ExAC (Lek et al., 2016) database with minor allele frequency of more than 5%; (6) Removing mutations mapped to non-coding regions such as introns, intergenic regions and non-coding RNAs; (7) Removing RNA editing sites listed in the DARNED (Kiran and Baranov, 2010) and RADAR (Ramaswami and Li, 2014) databases; (8) In cases where multiple reads that support a variant were aligned to the exact same positions, we kept only a single read; (9) Removing mutations that may be caused by sequencing leakage errors which are identified as alternate bases that have at least 3 bases that match the alternate base in a ±3 base window around the variant base; (10) Removing sites falling into pseudogenes or IgG genes which have noisy alignments. The tool is available upon request.

### Signature analysis

To identify mutational signatures we used our SignatureAnalyzer tool (Kim et al., 2016). Each mutation was associated with a signature if its likelihood of association to the signature was >0.75. We assigned a sample to a specific signature if more than 75% of its mutations were expected to arise from that signature (determined by the *H* matrix). To examine whether mutational signatures can be detected in data with a low number of mutations, we used the TCGA data analyzed in this study, in which 7 different mutational signatures were detected. We selected a subset of the called mutations, such that the distribution of mutation number per sample is identical to that found in GTEx and ran the SignatureAnalyzer tool.

### GTEx samples

We used all GTEx RNA-seq data that had matched WES data from the analysis freeze of release V7 (dbGaP accession phs000424.v7.p2), from all primary tissues except for blood. We removed samples had a RNA Integrity Number (RIN) smaller than 5.9. The full list of analyzed samples is in Supplementary Table 16. When applying RNA-MuTect to the GTEx samples we first observed a significant strand bias with an enrichment of C>A mutations on the non-transcribed strand (Haradhvala et al., 2016) (or G>T mutations on the transcribed strand) (Binomial P-value < 4.9×10^−324^, **Figure S11**). These mutations are likely to arise due to oxidation of guanines which have a known signature (8-oxoG) that results with G>T mutations. This finding implies that the oxidized guanine was on the transcribed RNA alone and not on the DNA. This phenomenon was not observed when analyzing TCGA RNA-seq data, and hence were likely generated as part of the GTEx library preparation process. We therefore removed all of these mutations from further analysis of the GTEx data.

### Identifying individuals with a significant difference in the total number of mutations

We first split the data into different categories, such that each category is defined by the combination of tissue site (*n* = 50), age (higher or lower than 45, *n* = 2), race (*n* = 4) and gender (*n* = 2), yielding overall 800 categories. Then, within each category *c*, we ranked the individuals associated with it by their number of mutations. This was done by first equally splitting the interval *x*= [0 1] into the number of individuals associated with a category (*n*_*c*_). We then considered the middle point of each segment in this interval 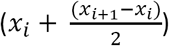 and assigned each individual with a value *r* in *x* according to its descending rank (lower numbers were assigned a higher rank). In cases where there were several individuals with the same number of mutations, we assigned them with the average value of their ranks, as described above. These ranks were then transformed into a score, *s*, such that 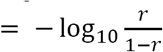. Finally, we summed up these scores associated with all the tissues of a given individual, and calculated the significance of the total score based on 10,000 permutations in which we shuffled the number of mutations within each category. We computed a two-sided empirical P-value, examining if there are individuals with a significantly higher or lower score. Empirical P-values were then corrected for multiple hypothesis using Benjamini-Hochberg FDR procedure (Benjamini and Hochberg, 1995) and considered significant if q-value ≤ 0.1.

### Studying the relation between expected number of mutation and expression level

For each gene *i* and sample *j*, we used a binomial model to compute the power to detect >=4 reads supporting the alternate allele, given the median allele fraction of 0.05 and the coverage per base. The power and coverage per gene in each sample was then computed as the mean power and coverage over the gene’s bases. We then split the data into 5 bins according to gene coverage, and computed the underlying number of mutations in each bin, as the overall number of mutations divided by the sum of power in that bin. This was done using the bootstrapping and subsampling with repeats method.

### Identification of highly expressed genes

To identify the set of genes that are highly expressed in the outlier vagina sample, we performed the following analysis: We first considered only genes that are expressed in this tissue (log_2_(TPM+1)>1); Then, we performed a one sided T-test to check if the expression value of each expressed gene is significantly higher compared to all other vagina samples; Our final gene list of 103 genes included ones that had a Bonferroni-corrected P-value < 0.05 and a fold-change > 2; This list was tested for GO enrichment using the GSEA tool (Subramanian et al., 2005).

### Association of number of mutations with proliferation index (*MKI67*)

To split all tissue types (that had more than 10 samples) into low and high mutation number, we used as a threshold the median value of the average number of mutations per tissue, such that half of the tissue types are considered with a high number of mutations and half with a low number of mutations. We considered tissues with a higher proliferation rate as those having (log_2_(TPM+1)>1) in more than 50% of their samples.

### Enrichment for driver genes

To compute enrichment per tissue for cancer genes we used the Cancer Gene Census (CGC) genes and focused only on genes that harbor missense or nonsense driver mutations (*n* = 249). We then computed a hypergeometric P-value, testing for a significant overlap between CGC genes and non-silent mutated genes in each tissue, out of all protein coding genes. To control for gene length and depth of coverage, we randomly selected 100 groups of *n* genes from each tissue that had a similar length and expression level to that of the tested CGC genes. Specifically, for each CGC gene we picked up to 10 most similar genes, in terms of expression level and length. In each iteration, we randomly selected one of these genes to replace by the corresponding CGC gene. We verified that all selected groups had a similar distribution as the CGC genes in the corresponding tissue (Wilcoxon P-value > 0.9 for both length and expression levels). We then computed an empirical P-value per tissue using the hypergeometric P-value as a test statistic, and controlled for multiple hypothesis testing using FDR across the tissues. Tissues significantly enriched with CGC genes were then selected as those with empirical q-value < 0.1.

### Hotspot and *TP53* analysis

Hotspot mutations were defined as mutations in cancer genes (CGC, Forbes et al., 2015) that appear more than three time in the TCGA consensus somatic mutation call set (doi:10.7303/syn7214402) (**Supplementary Table 11**). To test the allele fraction of *TP53* or hotspot mutations relative to all other mutations detected in that sample, we focused on 19 (out of 22) *TP53* mutations and on 22 (out of 30) hotspot mutations from samples that harbor overall ≥3 mutations. Then, for each mutation we replaced its allele fraction with its percentile within the sample and calculated how many of the mutations were above the 60^th^ percentile. To compute an empirical P-value for the hotspots we randomly drew 1000 groups of 22 mutations (one from each sample) and compared the number of mutations above the 60^th^ percentile in the randomizations compared to the observed. In addition, we performed another permutation test in which we control for gene coverage. This was done by randomly drawing groups of 22 mutations, such that for each gene we only select mutated genes with matching coverages (out of the closest 20 mutated genes). For both *TP53* and the hotspots, we applied another statistical test by randomly drawing one mutation from each of the samples associated with the tested mutations, and similarly examined how many of them were above the 60^th^ percentile. Empirical P-value was calculated based on a 1000 randomizations.

### Computing context dependent dN/dS

For each gene *i* we extracted the coding sequence of all of its possible transcripts. Per transcript *i*_*tr*_ and context *j* (*C*_*j*_) we computed a value 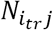 and 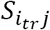 that represent the fraction of possible non-silent and silent mutation in context *j*, respectively. We then averaged these values for each gene *i* across its transcripts to obtain *N*_*ij*_ and *S*_*ij*_. Next, to obtain the overall chance of observing a non-silent mutation in gene *i*, we normalized these values using the 96 possible contexts as follows: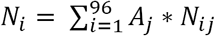 and 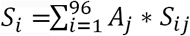, where *A*_*j*_ is a normalization factor that is proportional to the frequency of observing *C*_*j*_ in our data (the GTEx somatic mutations) and in gene *i*. We then used a binomial model to compute, for each gene *i*, the probability of observing at least *k* non-silent mutations (using *N*_*i*_), given a total number of *n* mutation in the gene. Finally, we compared sets of genes by applying Wilcoxon ranksum test to their corresponding distributions of p-values.

### MutSigCV for RNA-seq data

To apply MutSigCV (Lawrence et al., 2014) for RNA-seq data, we first generated a gene-level coverage model that reflects which bases are typically sufficiently covered in each gene using GTEx RNA-seq data. When applied to normal samples, and since the number of detected mutations is low, we focused the analysis on a combined list of 718 cancer genes from CGC and Lawrence *et al*. (Lawrence et al., 2014) (**Supplementary Table 13**). We estimated significance based on the overall burden of mutations (compared to the background model) as measured by *P*_*CV*_ in MutSigCV. We considered genes as significantly mutated if they had a FDR-corrected q-value < 0.05 and appeared as mutated 3 times or more. After a manual review we have decided to remove the NACA gene from the list. The association of this gene with cancer according to COSMIC is through a translocation with BCL6. As this is not the case in our data we believe that its significance might represent a false positive. To compute an empirical P-value for estimating the ranking of these genes’ allele fraction relative to others, we repeated the same analysis described for hotspots and *TP53* above.

### Variant detection in blood DNA

We focused on the set of 332 single nucleotide variants previously detected in the blood of healthy individuals (Jaiswal et al., 2014). We applied MuTect to obtain the reference and alternate allele counts of these mutations, in the blood of all individuals and their corresponding tissues. Following Jaiswal et al. (Jaiswal et al., 2014), we identified potential somatic mutations by identifying variants with allele fractions that are significantly lower than 0.5 (which is expected for heterozygous variants), using a binomial model given the coverage at the site. mutations with a Bonferroni-corrected P-value < 0.05, supported by at least 3 reads were considered as somatic blood mutations.

### Allelic imbalance analysis

Germline variants were detected in each individual using GATK HaplotypeCaller (DePristo et al., 2011). We then utilized MuTect (Cibulskis et al., 2013) in order to consistently count the number of reads supporting the reference and alternate alleles of the identified SNPs in the corresponding RNA-seq samples. We filtered out sites that (1) had an allele fraction (AF) smaller than 0.05 or greater than 0.95 in the normal DNA; (2) had a coverage of less than 10 reads in the RNA-seq. Then, for each SNP we considered the minimal value between *AF* and (1 - *AF*) and used this value to compute the mean AF for each chromosome arm, i.e. *mean*(min(*AF*, 1 - *AF*)) = *avg_min_AF*. Some of the RNA-seq samples were particularly noisy and were removed from the analysis. Specifically, we removed 329 (out of 6702) samples in which more than 15% of their RNA SNPs had an AF less than 0.3 or greater than 0.7. To identify significant allelic imbalance events, we fit a beta distribution per chromosome arm to the *avg_min_AF* across all samples that had at least 100 heterozygous sites. The fitted beta parameters were then used to compute a one-sided P-value per arm and per sample. P-values, for each chromosome arm, were corrected for multiple hypotheses using FDR to identify samples with significant imbalance (q-value < 0.05). To be even more conservative, we considered only significant chromosome arms that had the lowest *avg_min_AF* value compared to the other arms in the same sample, as computed by (min(*avg_min_AF*) - 3 \**std*(*avg_min_AF*)).

### Multiplex Fluidigm Access Array Validation by Resequencing for GTEx

Validation of mutations was performed by targeted resequencing using microfluidic PCR on the 48.48 Fluidigm Access system (Fluidigm, South San Francisco, CA) and the MiSeq sequencing system (Illumina, San Francisco, CA). 14 Samples were selected for validation based on the presence of mutations found by RNA sequencing. Target-specific primers were designed to flank sites of interest. PCR was performed on the Fluidigm Access Array according to the manufacturer’s instructions using the multi-plex protocol. The Access Array Integrated Fluidic Circuit (IFC) enabled parallel amplification of up to 48 unique samples per chip. The amplicon-tagging PCR, using tailed target-specific primers (tailed with adapter sequence), occurred on the microfluidic chip. This PCR product was then used as input for the molecular barcoding PCR, using primers containing sequence complementary to the target-specific primer tails, a molecular barcode, and flow cell attachment sequence (compatible with Illumina). This second PCR occurred within a 96-well plate on a standard thermocycler. The Bravo Automated Liquid Handler was used for multiplexing of primers, chip loading, PCR set-up and harvesting (Agilent Technologies, Lexington, MA). Indexed libraries (pools of amplicons) were quantified, and quality-checked using Caliper GX (Perkin Elmer, Boston, MA). These per-sample-amplicon-pools were then normalized based on concentration, and pooled into a single tube. Final amplicon library pools were quantified by qPCR using the Kapa Library Quantification Kit for NGS (Kapa Biosystems, Wilmington, MA), and sequenced on 2 MiSeqs, according to manufacturer’s protocol, using paired-end 150 bp sequencing reads.

## Code availability

The code of RNA-MuTect would be made available upon publication.

## Contributions

K.Y., P.P and G.G. conceived the idea. K.Y. and G.G designed the study. J.H, helped with the MutSig analysis. F.A., J.K., C.S., H.Z., D.L. and D.R. contributed code for the analysis. K.K. and P.B. reviewed the pathological samples. J.G. performed the Fluidigm experiments. R.F. generated the docker with the RNA-MuTect pipeline. X.L., E.L., N.S., A.S. and K.A helped with the interpretation of the data. K.Y. and G.G wrote the manuscript.

## Acknowledgements

We would like to thank Benjamin Ebert, Adam Bass, Todd R. Golub for helpful comments on the manuscript. We would also like to thank Julie Gastier-Foste and Erik Zmuda for helpful information regarding TCGA samples. K.Y was funded by the Broad-ISF postdoctoral fellowship and the Weizmann award for Women in Science. G.G was partially funded by the GTEx LDACC (HHSN268201000029C) and the Paul C. Zamecnick, MD, Chair in Oncology at MGH.

